# CARD8 inflammasome mediates pyroptosis of HIV-1-infected cells by sensing viral protease activity

**DOI:** 10.1101/2020.09.25.308734

**Authors:** Qiankun Wang, Hongbo Gao, Kolin M. Clark, Pengfei Tang, Gray H. Harlan, Sebla B. Kutluay, Carl J. DeSelm, Rachel M. Presti, Liang Shan

## Abstract

HIV-1 has high mutation rates and exists as mutant swarms in the host. Rapid evolution of HIV-1 allows the virus to outpace host immune system, leading to viral persistence. Novel approaches to target immutable components are needed to clear HIV-1 infection. Here we report a pattern-recognition receptor CARD8 that senses enzymatic activity of the HIV-1 protease, which is indispensable for the virus. All subtypes of HIV-1 can be sensed by CARD8 despite substantial viral diversity. HIV-1 evades CARD8 sensing because the viral protease remains inactive in infected cells prior to viral budding. Induction of premature intracellular activation of the viral protease triggers CARD8 inflammasome-mediated pyroptosis of HIV-1-infected cells. This strategy leads to clearance of latent HIV-1 in patient CD4^+^ T cells after virus reactivation. Taken together, our study identifies CARD8 as an inflammasome sensor of HIV-1 that holds promise as a strategy for clearance of persistent HIV-1 infection.

## Introduction

The adaptive immune system recognizes cognate epitopes based on amino acid sequences and associated conformations that are subject to mutation. Retroviruses mutate at very high rates largely due to the lack of proofreading activity of their reverse transcriptase (*1*). The within-host diversity of human immunodeficiency virus type 1 (HIV-1) allows rapid selection of antibody and T cell escape variants (*2-7*). In patients who received antiretroviral therapy (ART), HIV-1 persists in a latent form primarily in quiescent CD4^+^ T cells (*8-10*) and possibly tissue macrophages(*11*). Immune escape variants achieved in the latent viral reservoirs present one of the major obstacles to HIV-1 eradication (*12-15*). In addition, host cells persistently infected with HIV-1 are long-lived and resistant to virus- or immune-mediated apoptotic cell death (*16, 17*). In this study, we evaluated whether immune system could: 1) sense the functions of an essential HIV-1 protein including those with viral enzymatic activities, which are highly immutable; 2) mediate robust target cell killing independent of apoptosis. The host immune system utilizes germline-encoded pattern-recognition receptors (PRRs) to detect microbial products. A set of intracellular PRRs characterized by the presence of a caspase recruitment domain (CARD) or a pyrin (PYD) domain co-oligomerize with pro-caspase-1 and form high-molecular weight inflammasome complexes upon sensing of their cognate ligand (*18*). Inflammasome assembly in response to cytoplasmic microbial or danger signals leads to caspase-1 (CASP1) activation and pyroptosis, an inflammatory form of programmed cell death. There is no evidence that HIV-1 infection can be sensed by innate sensors associated with inflammasome activation in CD4^+^ T cells, with the exception of the bystander effect triggered by abortive HIV-1 transcripts (*19*). The bystander cell death mechanism only applies to quiescent lymphoid-resident CD4^+^ T cells that are non-permissive to HIV-1. It does not apply to the cells productively or latently infected with HIV-1, and thus has no impact on persistently infected cells. In the present study, we aimed to identify the inflammasome sensor(s) that recognizes intracellular HIV-1 activities.

## Results

### CARD8 senses HIV-1 protease activity

The caspase recruitment domain-containing protein 8 (CARD8) has been implicated in inflammasome activation and pyroptosis of CD4^+^ T cells and macrophages (*20-22*). A key question is whether CARD8 is an inflammasome sensor, and if so, what pathogens physiologically activate it. The C-terminus of CARD8 protein contains a ‘function-to-find’ domain (FIIND) followed by a CARD domain. Although the mouse genome does not encode CARD8, murine NLRP1b C-terminus also contains the FIIND-CARD domain. NLRP1b can be activated via direct proteolysis of its N-terminus by *Bacillus anthracis* lethal factor protease (LF) (*23*). Briefly, N-terminal cleavage of murine NLRP1b creates a neo-N-terminus which is ubiquitylated by the N-end rule pathway and targeted for proteasome degradation (*24, 25*). Due to the break in the polypeptide chain by FIIND auto-processing, the C-terminal bioactive subunit is liberated from the proteasome complex and can initiate CASP1-dependent inflammasome assembly (*26, 27*). Since human CARD8 shares structural similarity with murine NLRP1b and also undergoes autoprocessing (*28*), we hypothesized that CARD8 also might sense microbial protease activity (Fig. 1A). To determine whether CARD8 is a sensor for HIV-1 protease, HEK293T cells were co-transfected with an HIV-1 reporter vector pNL4-3 and human CARD8 vector with an N-terminal HA tag. In cells transfected with CARD8 alone, both the full length and the N-terminal fragment due to autoprocessing were detected by a CARD8 N-terminus antibody or an HA antibody. The viral plasmid pNL4-3 expresses all viral proteins except the envelope protein. The HIV-1 gene *pol* encodes three viral enzymes include protease, reverse transcriptase and integrase. In HIV-1-infected cells, the viral protease is embedded in the Gag-pol precursor protein and remains inactive. The viral protease activation requires dimerization of Gag-Pol polyprotein, which occurs during or soon after the viral budding process. Overexpression of HIV-1 Gag-Pol polyprotein in transfection systems resulted in its intracellular dimerization and premature protease activation (*29*). When the HIV-1 vector was co-transfected, a smaller N-terminal fragment was detected by the CARD8 N-terminus antibody as well as an HA antibody (Fig. 1B), suggesting that the HA-tagged N-terminus was cleaved by HIV-1 protease. In contrast, CARD8 was not cleaved by the viral vector when the viral protease was inactivated by a single mutation (D25H), as evidenced by the lack of HIV-1 Gag (p55) cleavage. To evaluate whether other viral proteins are required for CARD8 cleavage, we introduced various mutations to the viral vector (fig. S1). Cleavage of CARD8 was not affected when mutations were introduced to any viral genes except *pol*, but was blocked by an HIV-1 protease inhibitor lopinavir (LPV) (Fig. 1C). HIV-1-specific non-nucleoside reverse transcriptase inhibitors (NNRTI) such as efavirenz (EFV) and rilpivirine (RPV) but not nevirapine (NVP) can bind to HIV-1 Pol and enhance intracellular Gag-Pol polyprotein dimerization, which causes premature protease activation (*30*). As expected, addition of RPV further enhanced the HIV-1 protease-mediated cleavage of CARD8 and HIV-1 Gag (Fig. 1D). As initial study reported (*20*), the level of endogenous expression of CARD8 in HEK293T cells is low, not sufficient for the detection of N-terminal cleavage (fig. S2A). However, it might be sufficient to trigger the downstream signaling cascade upon HIV-1 protease cleavage. In cells transfected with CASP1, pro-IL1β and HIV-1 vectors with a functional viral protease, pro-IL-1β was processed into mature IL-1β (p17) without the need to overexpress CARD8 (Fig. 1E). As expected, IL-1β processing was enhanced by RPV, but blocked by LPV. To test whether proteasome degradation was required, we added proteasome inhibitors MG132 or bortezomib (bort) together with RPV into transfected cells. The baseline level of IL-1β-p17 was not affected by proteasome inhibitors because the inhibitors were added 24 hours post transfection. RPV-induced processing of pro-IL-1β was blocked by bort and MG132, while adding proteasome inhibitors did not affect HIV-1 Gag cleavage (Fig. 1F), which ruled out the possibility that MG132 and bort directly affected HIV-1 protease activity. We also generated CARD8 knockout cells to confirm the indispensable role of CARD8 in viral protease mediated inflammasome activation (data in Fig. 2 and 3). In contrast to transfection, infection of HEK293T cells with VSV-G pseudotyped HIV-1 reporter virus did not result in CARD8 cleavage due to the lack of protease activation. Addition of RPV was sufficient to activate HIV-1 protease and cleave CARD8 N-terminus, which was blocked by LPV (Fig. 1G). In summary, we demonstrated that HIV-1 protease could degrade the N-terminus of CARD8 and that activation of the viral protease could lead to proteasome/CASP1-dependent inflammasome activation in the HEK293T transfection system.

**Fig. 1.**
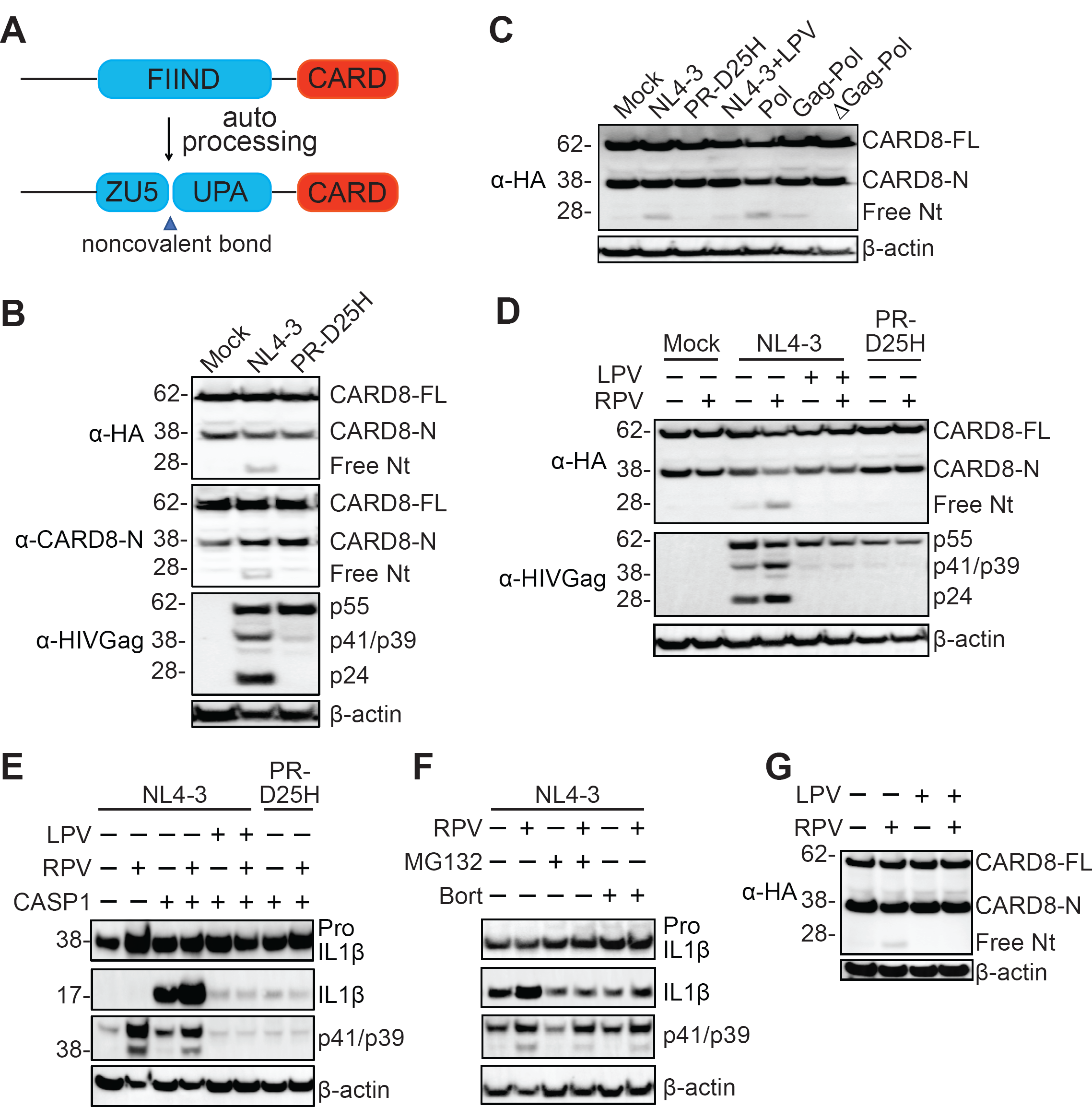
HIV-1 protease mediates CARD8 cleavage. (**A**) Domain architecture of the CARD8 protein. (**B**) HIV-1 protease cleaves the N-terminus of CARD8. HEK293T cells were transfected with plasmids encoding HA-CARD8 (100ng), together with either the pNL4-3 (1μg) or Pro-D25H (1μg). Cells were collected 24 hours after transfection. Anti-HA, anti-CARD8-N and anti-p24 antibodies were used sequentially on the same blot. (**C**) HIV-1 protease is necessary and sufficient to cleave CARD8. HEK293T cells were transfected with constructs encoding CARD8 (100ng) together with indicated viral plasmids (1μg). Cells were collected 24 hours after transfection. (**D**) RPV enhances HIV-1 protease-mediated cleavage of CARD8. HEK293T cells were transiently transfected with HA-CARD8 (100ng) and indicated viral plasmids (1μg). DMSO or RPV was added 24 hours post transfection. Cell lysates were collected 6 hours after RPV treatment. (**E and F**) HIV-1 protease triggers CASP1-dependent pro-IL-1β processing. HEK293T cells were co-transfected with plasmids encoding CASP1 (2ng), pro-IL-1β (200ng) and an HIV-1 plasmid (1μg). After 24 hours, cells were treated with indicated drugs for another 6 hours. (**G**) RPV induces HIV-1 protease-dependent cleavage of CARD8 in infected cells. HEK293T cells were infected with VSV-G-pseudotyped HIV-1 reporter virus. Infected cells were then transfected with HA-CARD8 (100ng). DMSO or RPV was added 24 hours post transfection. Cell lysates were collected 6 hours after RPV treatment. In **B**-**F**, cell lysates were evaluated by immunoblotting. CARD8-FL, full-length CARD8; CARD8-N, N terminal CARD8; free Nt, freed N terminus. Data are representative of three or more independent experiments.

**Fig. 2.**
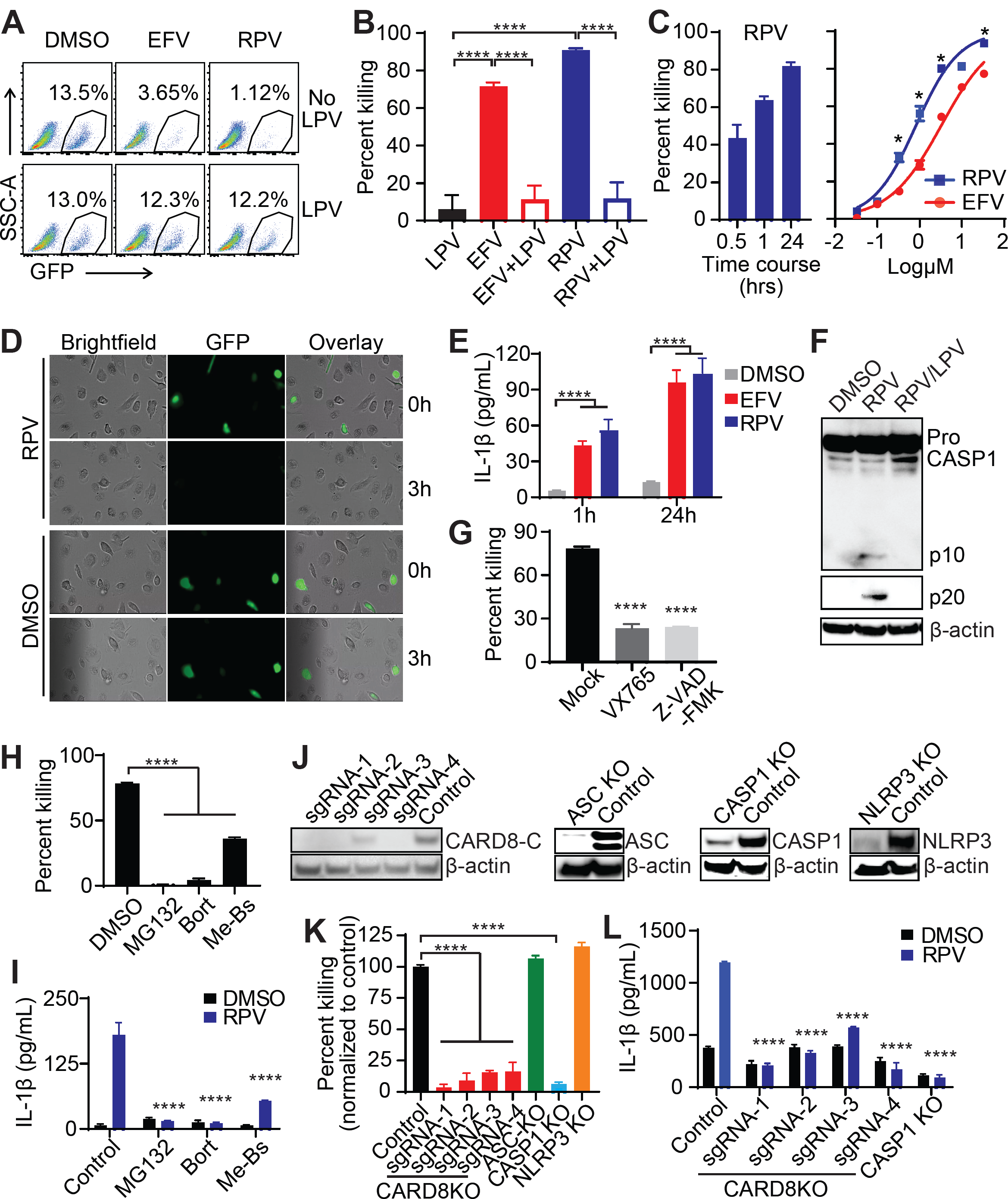
HIV-1 protease triggers CARD8-dependent pyroptosis of infected macrophages. (**A to E**) HIV-1 protease activation by NNRTIs induced rapid pyroptosis of infected monocytes derived macrophages (MDMs). MDMs were infected with HIV_NL4-3/BaL_. At day 4, Raltegravir (RAL) and T-20 were added to block new infection. Cells were then treated with RPV, EFV, LPV, or combinations for up to 24 hours. In **A to C**, GFP^+^ cells were detected by flow cytometry. In **D**, images of infected MDMs were taken using a Cytation 5 Imaging Multi-Mode Reader (Biotek). In **E**, culture supernatant was collected for IL-1β ELISA. (**F** and **G**) Pyroptosis of HIV-1-infected MDMs is CASP1-dependent. MDMs were infected and treated as described above. In **F**, immunoblot analysis of pro-(p45) and cleaved CASP1 (p10 and p20) in infected MDMs after RPV treatment for 1 hour. In **G**, infected MDMs were pre-treated with VX-765 (100μM) or Z-VAD-FMK (100μM) for 3 hours and then treated with RPV for 4 hours before flow cytometry analysis. (**H** and **I**) HIV-1 protease mediated inflammasome activation is proteasome-dependent. MDMs were infected and treated as described above. Infected MDMs were pretreated with proteasome inhibitors MG132, Bort, or Me-Bs for 30 minutes and then treated with RPV for 4 hours. In **H**, GFP expression was analyzed by flow cytometry. In **I**, culture supernatant was collected for the detection of IL-1β by ELISA. (**J** to **I**) The CARD8 inflammasome is required for pyroptosis of HIV-1-infected macrophages. (**J**) knockout of CARD8, ASC, CASP1 or NLRP3 in THP-1 cells was confirmed by immunoblotting. Knockout or control THP-1 cells were infected with VSV-G pseudotyped HIV-1 reporter virus NL4-3-Pol. 3 days after infection, cells were pre-treated with LPS (100ng/ml) for 3 hours before RPV treatment. In **K**, GFP expression was analyzed by flow cytometry 24 hours post RPV treatment; Data were normalized to the control group. In **L**, culture supernatant was collected 48 hours post RPV treatment for IL-1β detection. In **B, G, H** and **K**, *P* values were calculated using one-way ANOVA and Turkey multiple comparison tests. In **C, E, I** and **L** *P* values were calculated using two-way ANOVA and Turkey multiple comparison tests. *p < 0.05, ****p < 0.0001. In each bar graph, n≥3. Error bars show mean values with SEM. Data are representative of three or more independent experiments.

**Fig. 3.**
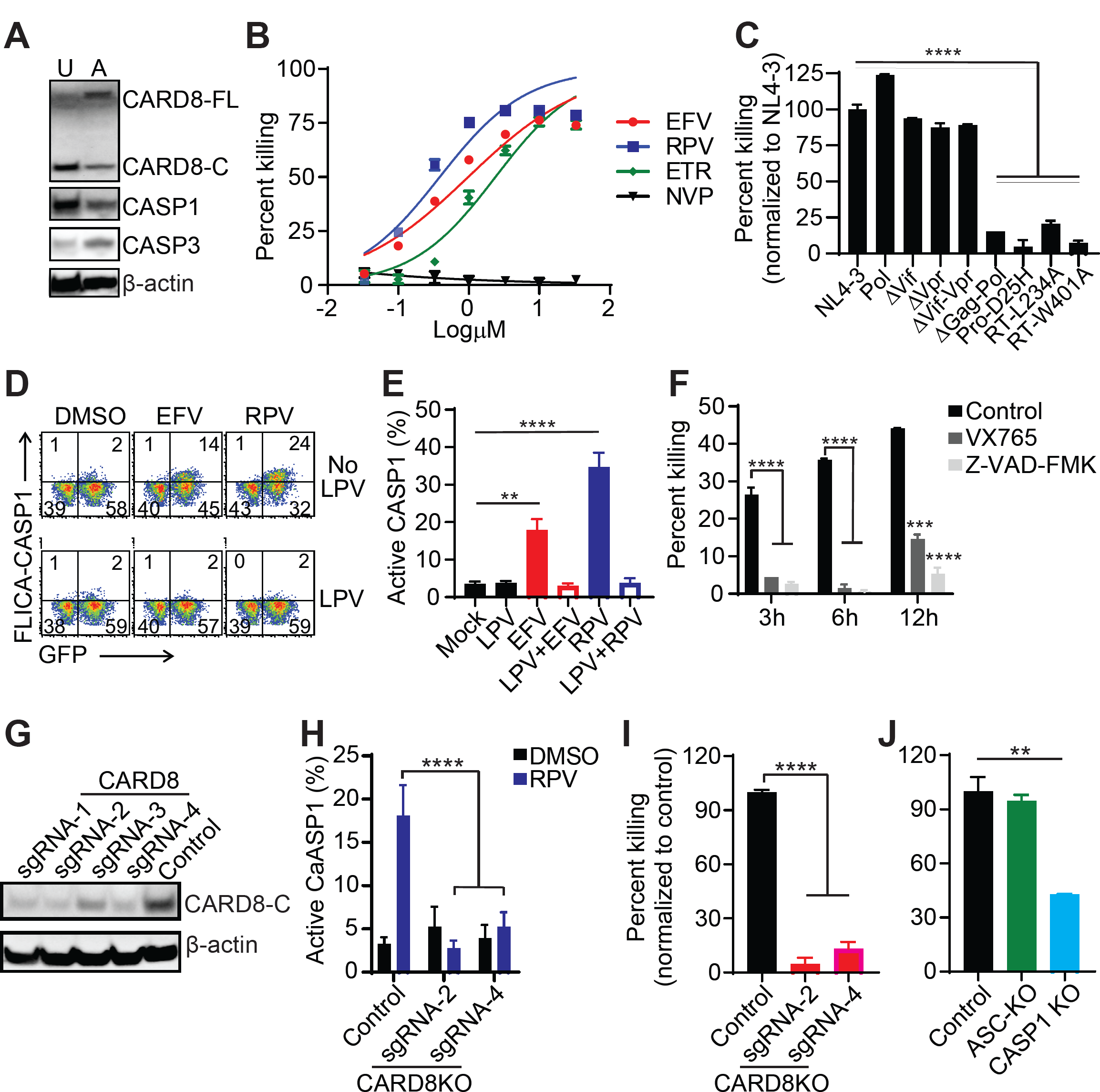
HIV-1 Protease induces CARD8 inflammasome activation and subsequent cell death in CD4^+^ T cells. (**A**) Analysis of CARD8, Caspase 1 and Caspase 3 expression level in unstimulated (U) and activated (A) primary CD4^+^ T cells by immunoblotting. (**B** and **C**) HIV-1 protease activation by NNRTIs leads to killing of infected primary CD4^+^ T cells. Activated CD4^+^ T cells were used for viral infection. In **B**, cells were infected with HIV-1 reporter virus NL4-3-Pol. Various NNRTIs at indicated concentration were added 3 days post infection. In **C**, cells were infected with different HIV-1 reporter viruses for 3 days and treated with RPV. Cells were analyzed for GFP expression by flow cytometry 48 hours post NNRTI treatment. (**D** to **F**) HIV-1 protease activation by NNRTIs induced CASP1 activation in infected primary CD4^+^ T cells. Activated primary CD4^+^ T cells infected with the NL4-3-Pol. In **D** and **E**, infected cells were treated with EFV, RPV, LPV and combinations for 3 hours before FLICA660 Caspase 1 staining for active CASP1 detection. In **F**, infected cells were pre-treated with CASP1 inhibitor VX765 (50μM) and Pan-Caspase inhibitor Z-VAD-FMK (50μM) for 3 hours before adding RPV. GFP expression was measured by flow cytometry at indicated time. (**G** to **J**) The CARD8 inflammasome is required for pyroptosis of HIV-1-infected primary CD4^+^ T cells. In **G**, knockout of CARD8 in primary CD4^+^ T cells was confirmed by immunoblotting. **H** to **J**, CARD8, ASC or CASP1 knockout primary CD4^+^ T cells were co-stimulated and then infected with NL4-3-Pol. At day 3 post infection, cells were treated with RPV. In **H**, FLICA660 Caspase 1 was used for staining of active CASP1 at 3 hours post RPV treatment. In **I** and **J**, GFP expression was measure by flow cytometry 24 hours after RPV treatment. In **C, E, I** and **J**, *P* values were calculated using one-way ANOVA and Turkey multiple comparison tests. In **F** and **H**, *P* values were calculated using two-way ANOVA and Turkey multiple comparison tests. **p < 0.01, ***p < 0.001, and ****p < 0.0001. In each bar graph, n≥3. Error bars show mean values with SEM. Data are representative of five or more independent experiments.

### HIV-1 triggers CARD8-depedent pyroptosis of infected macrophages

CARD8 is expressed in HIV-1 target cells including primary CD4^+^ T cells and macrophages (fig. S2A). The key question was whether induction of premature intracellular activation of HIV-1 protease could trigger CARD8 inflammasome activation and pyroptosis of infected cells. When treated with EFV or RPV, HIV-1-infected macrophages (GFP^+^) rapidly underwent pyroptotic cell death as evidenced by membrane swelling and rupture (Fig. 2, A to D and Video S1), as well as secretion of IL-1β (Fig. 2E). The cell death triggered by EFV or RPV was rapid and dose-dependent, and completely blocked by LPV (Fig. 2, B and C). CASP1 activation was evidenced by detection of its active subunits p10 and p20 (Fig. 2F). In addition, we tested CASP1 specific inhibitor VX765 and a pan caspase inhibitor Z-VAD-FMK. While most of the infected macrophages were killed by RPV, blocking of CASP1 activity reduced the loss of infected cells to 20% (Fig. 2G). The N-end rule pathway mediates proteasome degradation of mouse NLRP1b (*31*). To test whether it was required for CARD8 activation, we added MG132 and bort or an N-end rule pathway inhibitor Bestatin methyl ester (Me-Bs) together with RPV to HIV-1-infected MDMs. MG132, bort and Me-Bs effectively blocked RPV-mediated killing of infected macrophages and secretion of IL-1β (Fig. 2, H and I). As chemical inhibitors often have off-target effects, we confirmed that CARD8 inflammasome was responsible for HIV-1 sensing by generating CARD8, ASC, CASP1 and NLRP3 knockout THP-1 cells (Fig. 2J). Deletion of CARD8 or CASP1, but not ASC or NLRP3, inhibited HIV-1 protease-mediated pyroptosis and IL-1β secretion, which was consistent with the inhibitor experiments. Our finding also confirms studies showing that CARD8 can form an Asc-independent inflammasome complex (*32*).

### HIV-1 triggers CARD8-depedent pyroptosis of infected CD4^+^ T cells

Several studies have reported that NNRTIs could induce HIV-1 protease-dependent killing of HIV-1-infected CD4^+^ T cells, although the killing mechanism is unknown (*33-35*). We hypothesized that the cytolysis observed in those studies was due to NNRTI-triggered HIV-1 protease-mediated CARD8 inflammasome activation. Since resting CD4^+^ T cells are the most well characterized cellular reservoirs for HIV-1, we examined the expression levels of key components of the CARD8 inflammasome in different subsets of primary CD4^+^ T cells. CARD8 is expressed in both activated and unstimulated blood CD4^+^ T cells, as well as in memory and naïve CD4^+^ T cells in lymphoid tissues (Fig. 3A and fig. S2, A and B). Both unstimulated and activated CD4^+^ T cells were susceptible to HIV-1 protease triggered cell death upon NNRTI treatment (Fig. 3B and fig. S2C). NVP was the only NNRTI without killing effect because it could not drive Gag-Pol dimerization (*35*). Since several HIV-1 proteins can induce death of primary CD4^+^ T cells, we produced different reporter viruses carrying mutations in *vif, vpr, vpu, env* and *nef* by transfecting HEK293T cells with different viral plasmids (fig. S1). Reporter viruses without a functional protease (ΔGag-Pol and Pro-D25H) or deficient in Gag-Pol dimerization (RT-L234A and RT-W401A) were unable to trigger cell death. None of the other viral proteins were required for cell killing (Fig. 3C). In addition to the pseudotyped reporter virus, we also observed cell killing with a clinical isolate HIV_BaL_. Since HIV_BaL_ is a replication-competent virus, all classes of antiretroviral drugs blocked viral replication, while NNRTIs could further reduce viral infection by clearing cells already infected with HIV-1 (fig. S3). Next, we showed that CASP1 activation was induced by EFV or RPV, but blocked by LPV (Fig. 3, D and E). Both VX765 and Z-VAD-FMK blocked killing of HIV-1-infected CD4^+^ T cells (Fig. 3F). Similar to infected macrophages, HIV-1 protease triggered CASP1 activation and cell death was also blocked by MG132, bort and Me-Bs (fig. S4). Lastly, we generated CARD8 knockout primary CD4^+^ T cells (Fig. 3G) to directly assess whether killing of infected CD4^+^ T cells was CARD8-dependent. CARD8 knockout abrogated NNRTI-induced CASP1 activation and pyroptosis of HIV-1-infected primary CD4^+^ T cells (Fig. 3, H and I). Similarly, CASP1 knockout or knockdown in primary CD4^+^ T cells also conferred resistance to HIV-1 protease-mediated pyroptosis (Fig. 3J and fig. S5).

### Activation of CARD8 inflammasome clears latent HIV-1 in patient CD4^+^ T cells

To determine whether the viral protease function in activating the CARD8 inflammasome is conserved, we tested a panel of HIV-1 virus isolates from chronically infected individuals of subtypes A, B, C and D (*36*). Subtype B is the dominant subtype in Europe and North America, while A, C and D are more important worldwide. T-20 and Raltegravir (RAL) were used to completely block new infection (data not shown) but had no killing effect. Addition of EFV or RPV effectively cleared primary CD4^+^ T cells infected with all HIV-1 subtypes, and the killing efficiency was not correlated with viral replication fitness (Fig. 4A and fig. S6), suggesting that the enzymatic activity of HIV-1 protease with regard to CARD8 activation is well conserved across major HIV-1 subtypes. To test whether strategies involving targeted activation of CARD8 inflammasome could be used for the clearance of latent HIV-1, we obtained blood CD4^+^ T cells from patients under suppressive ART. As shown in Fig. 4B, we performed limiting dilution of patient CD4^+^ T cells and seeded the cells into culture plates for virus reactivation and RPV treatment prior to quantitative viral outgrowth assay (*37*). We included NVP in the control group because it is the only NNRTI that has no cell killing function. The control ARV combination containing T-20, RAL and NVP had no killing effect, and was sufficient to completely block viral spreading (fig. S7, A and B). We found that short-term treatment with ARV combinations was not cytotoxic (fig. S7C). The infectious units per million (IUPM) was calculated as previously described^29^. The median IUPM in control and RPV groups is 2.61 and 0.16, respectively, suggesting a rapid clearance of 93.9% of the latent HIV-1 reservoirs (Fig. 4C). In fact, 3 out of 8 patient samples had no detectable viral replication after RPV treatment. In our “shock and kill” assay, cells were only treated with RPV for the first 2-3 days. It is possible that the residual viruses in the RPV group came from delayed virus reactivation which occurred after removal of RPV.

**Fig. 4.**
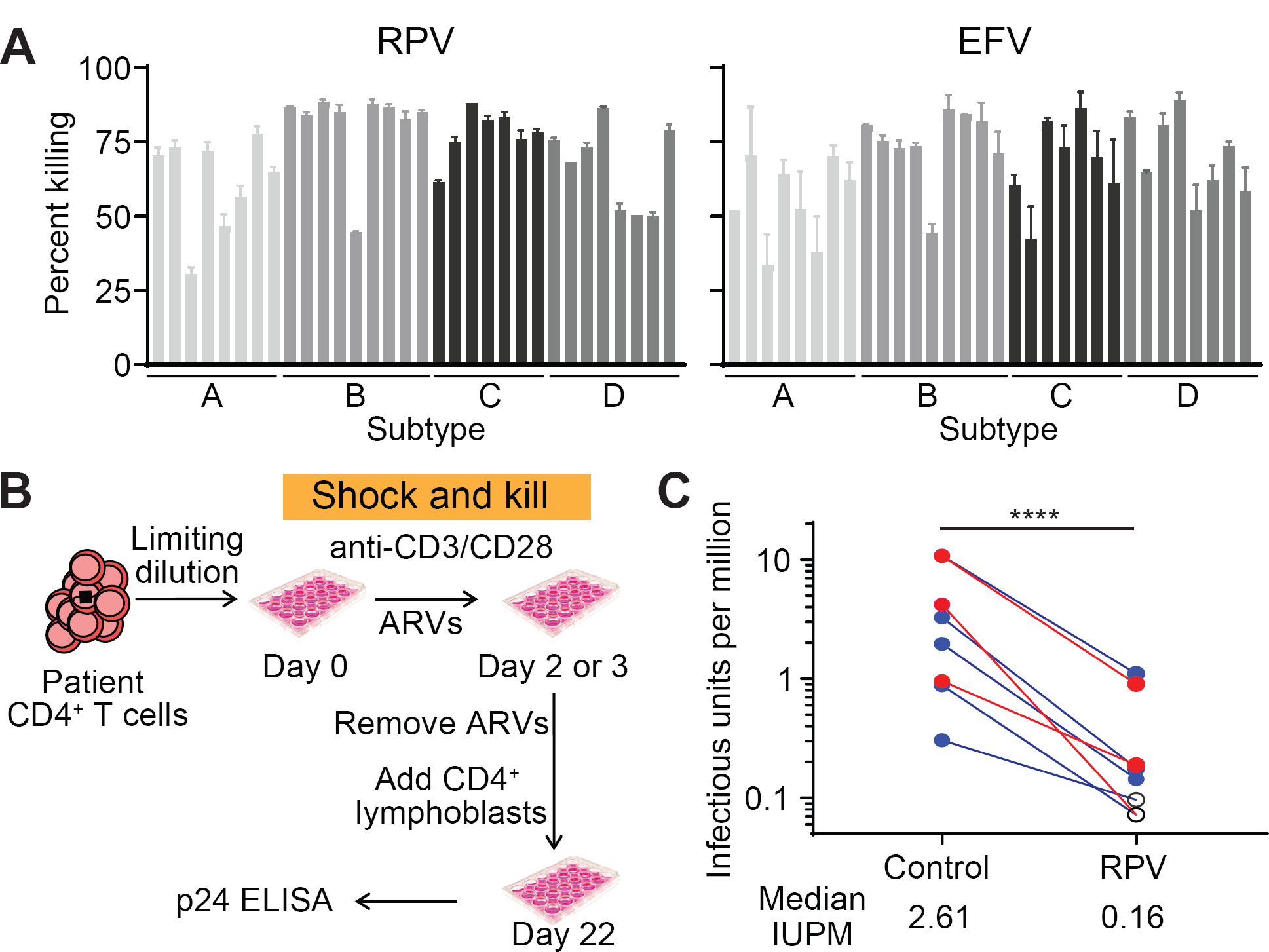
Induction of the CARD8 inflammasome activation clears infection of clinical HIV-1 isolates. (**A**) Killing of primary CD4^+^ T cells infected with different subtypes of clinical HIV-1 isolates. Activated CD4^+^ T cells were infected with a panel of international HIV-1 isolates. RAL (5μM) and T-20 (1μM) with or without EFV or RPV (5μM) were added on day 6 to block new infection. Viral infection was measured by intracellular p24 staining. N=3. Error bars show mean values with SEM. (**B**) scheme of shock and kill strategy using patient CD4^+^ T cells. Patient CD4^+^ T cells were seeded and stimulated to reactivate latent HIV-1 with the presence of antiretrovirals (ARVs). Control ARV combination includes NVP (5μM), RAL (10μM) and T-20 (10μM). RPV-containing combination includes RPV (5μM or 2.5μM), RAL (10μM) and T-20 (10μM). ARVs were removed at day 2 or 3. (**C**) Clearance of latent HIV-1 by RPV treatment. Quantitative viral outgrowth assay was performed to determine the frequency of latent HIV-1 after RPV or control treatment. Red: ARV for 2 days and RPV at 2.5μM. Blue: ARV for 3 days and RPV at 5μM. Open circle: no detectable HIV-1 infection by p24 ELISA. P value was calculated by ratio paired t test. *** P<0.001.

## Discussion

Due to rapid viral evolution, it is very difficult for the host immune system to control HIV-1 infection and clear residual viral reservoirs without targeting immutable components of the virus. In this study, we found that CARD8 is a sensor for HIV-1 protease activity to trigger inflammasome activation and pyroptosis of infected cells. Our study demonstrates that the human CARD8 and mouse NLRP1b inflammasomes share similar mechanism of activation, which involves their N-terminal cleavage by microbial proteases, followed by proteasome-mediated release of the bioactive C-terminal fragment to trigger inflammasome assembly and CASP1 activation. In HIV-1-infected cells, the CARD8 inflammasome cannot detect the virus because the viral protease remains inactive as a subunit of unprocessed viral Gag-Pol polyprotein. Surprisingly, some NNRTIs which have been used to treat HIV-1 infection for more than two decades can facilitate CARD8 sensing by mediating premature intracellular activation of HIV-1 protease. NNRTI-containing treatment regimens cannot eliminate HIV-1 infection in patients likely because the viral latent reservoirs are rapidly established prior to treatment initiation. Inclusion of NNRTIs in HIV-1 cure strategies should facilitate the elimination of infected cells after viral latency reversal. Intriguingly, CARD8 is preferentially and highly expressed in blood and lymphoid tissues (*38*) as well as in many hematopoietic-derived cells (*20-22*), suggesting that targeting the CARD8 inflammasome should be effective in lymphoid tissues, the most important anatomical sites for persistent HIV-1 infection. It is worth noting that the cell-killing IC_50_ of EFV and RPV is around 1-2µM (Fig. 2C and 3B), which is about 100-fold higher than the infection-blocking IC_50_. Although the plasma EFV concentration in patients receiving EFV-containing regiments (1-4µg/ml, or 3-12µM) is within the therapeutic range for cell killing, this strategy is unlikely to be effective in tissues with markedly lower drug concentration such as central nervous system. In addition, mutations in the HIV-1 Pol that confer resistance to NNRTIs also abrogate NNRTI-triggered cell killing (*35*) likely because the resistant viral variants can avoid drug binding. Therefore, the identification of new and more potent chemical compounds that promote Gag-Pol dimerization regardless of viral inhibition are warranted. Taken together, we discovered a mechanism of innate sensing of HIV-1 infection that has immediate implications for a strategy for HIV-1 cure.

## Materials and Methods

### Study participants

HIV-1-infected adult patients were recruited at the Infectious Diseases Clinic at Barnes-Jewish Hospital. Both male and female patients were included. All patients were on ART and had undetectable plasma HIV-1 RNA (< 20 copies per ml) for at least 6 months before blood collection. This study was approved by Washington University School of Medicine Internal Review Board. All study participants were provided with written informed consent. To obtain blood CD4^+^ T cells and monocytes for *in vitro* studies, anonymous peripheral blood samples were acquired from the Mississippi Valley Regional Blood Center as waste cellular products. Human tonsil biopsies with no identifiers attached were collected from elective tonsillectomies from Children’s Hospital in Saint Louis.

### Plasmids

The plasmids for production of replication-defective and replication-competent HIV-1 reporter viruses were listed in fig. S1. To obtain plasmids for replication-defective viruses, mutations were introduced into the pNL4-3-ΔEnv-EGFP vector (AIDS Reagent Program #11100), which has enhanced green fluorescent protein EGFP coding gene inserted into *env*. The plasmid pNL4-3/BaL was used for production of replication competent HIV-1 reporter virus HIV_NL4-3/BaL_. pNL4-3/BaL was modified using pNL4-3 with EGFP coding gene inserted into *nef* and *env* from the HIV_BaL_. The expression vector for HA-tagged CARD8 was constructed by RT-PCR to obtain cDNA fragment from primary CD4^+^ T cells and then subcloned it into the pcDNA3.1 vector. The CASP1 expression vector was from Addgene (#41552). The IL-1β (HG10139-CM) expression plasmid was purchased from Sino Biological. L40C-CRISPR.EFS.PAC (Addgene #89393) and SGL40C-H1.EFS.RFP657 (Addgene #69148) vectors were used to generate lentivirus vectors for sgRNA delivery. CRISPR/Cas9 target sites were selected using the CCTop selection tool. pLKO.1puro (Addgene #8453) was used to generate lentivirus vectors for gene-specific knockdown.

### Cell culture

HEK293T and THP-1 cells were obtained from ATCC and cultured in DMEM or RPMI 1640 medium supplemented with 10% fetal bovine serum (FBS), 1 U/ml penicillin and 100 mg/ml streptomycin (Gibco). Human CD14^+^ cells were isolated from healthy donor PBMCs using MojoSort Human CD14 Selection Kit (Biolegend #480026), and were cultured with 50 ng/ml recombinant human GM-CSF (BioLegend #572904) and 50 ng/ml recombinant human M-CSF (BioLegend #574806) for 6 days to obtain monocytes derived macrophages. CD4^+^ T cells from human tonsil biopsies were purified by sorting. Blood CD4^+^ T cells were isolated from healthy donor or patient PBMCs using a human CD4^+^ T cell isolation kit (BioLegend #480010). Purified CD4^+^ T cells were used without stimulation, or co-stimulated with plate-bound CD3 (Biolegend #300333) and soluble CD28 (Biolegend #302943) antibodies with the presence of 20 ng/μl IL-2 (Biolegend #589106) for 3 days.

### Antibodies and other staining reagents

Antibodies for N-terminal CARD8 (#ab194585), C-terminal CARD8 (#ab24186) and CASP1 p10 (#ab179515) were purchased from Abcam. The antibody for CARD8 C-terminus was used to detect endogenous CARD8 expression because of its high specificity. The antibody for CARD8 N-terminus was only used to detect the full-length and the cleaved N-terminus CARD8 in transfected HEK293T cells. Antibodies for CASP1 p20 (#4199), ASC (#13833), Caspase 3 (#9662), NLRP3 (#13158), Pro-IL-1β (#12703) and Cleaved IL-1β (#83186) were purchased from Cell Signaling Technology. Antibody for HA (#901502) was purchased from BioLegend. Antibody for beta-Actin (#MA1-140) and the secondary antibodies including horseradish peroxidase (HRP)-conjugated goat anti-mouse IgG (#31439) and HRP-conjugated goat anti-human IgG (#31410) were purchased from Invitrogen. The secondary antibody for HRP-conjugated goat Anti-rabbit IgG (#12-348) was from Sigma-Aldrich. HIV-1 p24 antibody (#530) for immunoblotting was obtained from NIH AIDS Reagent Program. HIV-1 p24-PE antibody (KC57-RD1, #6604667) for intracellular staining was purchased from Beckman Coulter. To sort T cell subsets from human tonsil biopsies, CD3 (Clone #HIT3a), CD4 (Clone OKT4), CD8 (Clone HIT8a), CD45RO (Clone # UCHL1) and CCR7 (Clone G043H7) were purchased from Biolegend. The FLICA660 Caspase1 staining reagents were purchased from ImmunoChemistry Technologies (#9122). Annexin V/7-AAD staining kit was purchased from Biolegend (#640930).

### Preparation of HIV-1 and lentivirus stocks

The replication-defective HIV-1 reporter viruses were prepared by co-transfecting HEK293T cells with viral vectors, a packaging vector pC-Help, and pVSV-G (Addgene #8454) or the NL4-3 envelop expressing plasmid. The replication-competent HIV-1 reporter viruses HIV_NL4-3/BaL_ was prepared by transfecting HEK293T with pNL4-3/BaL. To outgrow clinical HIV-1 isolates, international HIV-1 isolates (AIDS reagent program #11412) were used to infect CD8-depleted PHA-stimulated PBMCs. Culture supernatant was collected 6-9 days post infection. The knockdown or knockout lentiviruses were also produced in HEK293T cells by co-transfecting pVSV-G, psPAX2 (Addgene #12260), and sgRNA or shRNA lentiviral construct using Lipofectamine 2000 (Thermo Fisher). Lenti-X Concentrator (TaKaRa #631232) was used to obtain concentrated viral stock.

### Immunoblotting

4.0 × 10^5^ HEK293T cells were seeded in 12-well plates and cultured overnight before transfection using Lipofectamine 2000 (Thermo Fisher Scientific, 11668019). Lysates from 1.0 × 10^6^ cells were collected for protein expression analysis by immunoblotting. Cells were resuspended in RIPA buffer (Cell Signaling Technology #9806) with protease inhibitors (Thermo Scientific #78430). Protein concentrations were determined using the BCA protein assay kit (Thermo Scientific #A53225). Protein samples were separated by SDS–PAGE (Invitrogen #NW04125BOX), immunoblotted and visualized using ChemiDoc Imaging System (Bio-Rad). The CARD8 C-terminal antibody was used to detect endogenous CARD8 in THP-1 and primary CD4^+^ T cells. The CARD8 N-terminal antibody was used to detect N-terminal cleavage of CARD8 in transfected HEK293T cells.

### Gene transcription analysis

To determine gene transcription levels, total cellular RNA was extracted by Direct-zol RNA Kits (Zymo Research #R2071), and then reverse transcribed into cDNA (SuperScript™ III, ThermoFisher). Pre-designed gene-specific TaqMan assays were purchased for quantitative PCR. The transcription level of a housekeeping gene *polr2a* was measured for data normalization.

### Generation of THP-1 or primary CD4^+^ T cells with gene knockout or knockdown

The sgRNA and shRNA sequences were listed in fig. S1. All plasmids were verified by sequencing. For CARD8-KO, four sgRNAs were tested. The lentivirus vector for ASC, CASP1 or NLRP3 knockout carried two sgRNAs under the control of U6 and H1 promoter, respectively. THP-1 cells or activated primary CD4^+^ T cells were transduced with lentivirus carrying sgRNA or shRNA by spin inoculation for 2 hours at 1,200g at 25 °C with 8 μg/ml Polybrene. Transduced cells were then selected with puromycin (1μg/ml) for 5-7 days. For primary CD4^+^ T cells, after selection by puromycin, cells were re-stimulated with plate-bound CD3 and soluble CD28 antibodies with the presence of 20 ng/μl IL-2 (Biolegend #589106) for 3 days before infection with HIV-1 reporter viruses. Immunoblotting was performed to confirm knockout or knockdown efficiency.

### HIV-1 infection and drug treatment

HIV-1 infection was performed by spin inoculation for 2 hours at 1,200g at 25 °C. The following antiretrovirals (ARVs) were obtained from the NIH AIDS Research and Reference Reagent Program: Rilpivirine (RPV), Efavirenz (EFV), Lopinavir (LPV), Etravirine (ETR), Nevirapine (NVP), Maraviroc (MAV), T-20, Tenofovir (TFV) and Raltegravir (RAL). NNRTIs alone or in combination was added to HIV-1-infected cells 3-4 days post infection. ARVs were always used at 5μM, unless noted otherwise. CASP1 inhibitor VX765 was purchased from InvivoGene (#tlrl-vad). Pan-Caspase inhibitor Z-VAD-FMK was purchased from Selleckchem (#S2228). To block CASP1, infected cells were pre-treated with VX765 or Z-VAD-FMK for 3 hours before RPV or EFV treatment. Proteasome inhibitors MG132 (#10012628) and Bortezomib (Bort) (#10008822) were purchased from Cayman Chemical. Nonspecific aminopeptidase inhibitor Bestatin methyl ester (Me-Bs) was purchased from Abcam (#ab145909). To block proteasome degradation, transfected or infected cells were pre-treated with MG132 (10μM), Bort (5μM), or Me-Bs (20μM) for 30 minutes before RPV or EFV treatment. Cells treated with proteasome inhibitors were cultured for no more than 6 hours due to drug toxicity. IL-1β ELISA kit was purchased from BioLegend (#437004) for detection of IL-1β in culture supernatant. For GFP-reporter viruses, infection was determined by flow cytometry. For clinical isolates, intracellular HIV-p24 staining was performed using the Cytofix/CytopermTM kit (BD #554714). Flow cytometry was performed using BD LSRFortessa or BD accuri c6 plus. Flow cytometry data were analyzed by Flowjo software.

### Microscopy

HIV-1-infected MDMs were seeded in 96-well plates. Imaging started immediately after RPV was added to the cells. Images were acquired in random fields at 20x magnification in brightfield and GFP (469nm/525nm) channels using Cytation 5 Imaging Multi-Mode Reader (Biotek) at 3 minutes intervals for 6 hours. Conditions were maintained at 37 °C and 5% CO2 during live imaging. Images were processed and stitched using Gen5 software (Biotek). Representative time-lapse video depicts a single field over 6 hours.

### Quantitative viral outgrowth assay (QVOA)

QVOA was performed as previously described (*37*). Briefly, purified patient CD4^+^ T cells were with different antiretroviral combinations for Cells were plated at 1.0⨯10^6^, 0.5×10^6^, and 0.2⨯10^6^ cells/well before co-stimulation with anti-CD3/CD28 antibodies and 20 ng/μl IL-2 for 2 or 3 days. Cells were washed with PBS for 6 times to thoroughly remove ARVs and then co-cultured with CD8-depleted PHA-stimulated PBMCs from healthy donors. Culture supernatant was collected at 22 days for HIV-1 p24 ELISA (XpressBio #XB-1000). The frequency of p24 positive wells was used to calculate the estimate infection frequencies using a limiting dilution calculator (https://silicianolab.johnshopkins.edu/). Patient samples with undetectable latent HIV-1 (no p24^+^ wells) in the control group, or with NNRTI-resistant mutations were excluded in Fig. 4C.

### Statistical Analysis

Statistical analyses were performed using Prism 8 (GraphPad). The methods for statistical analysis were included in the figure legends. Error bars show mean values with SEM.

## Acknowledgements

We thank the volunteers for participating in this study. We thank Lisa Kessels, Michael Klebert, Alem Haile, Teresa Spitz, Timira Minor for patient recruitment. We thank Marina Cella for the tonsil biopsies. We thank Michael Diamond and Ruaidhri Jackson for comments and suggestions on this manuscript. The following reagents were obtained through the AIDS Research and Reference Reagent Program, Division of AIDS, NIAID, NIH: Rilpivirine, Efavirenz, Lopinavir, Etravirine, Nevirapine, Maraviroc, T-20, Tenofovir, Raltegravir, pNL4-3-ΔEnv-EGFP, international HIV-1 isolates, MOLT-4 CCR5^+^ Cells and HIV-1 p24 antibody. This work was supported by NIH grants R00AI125065 (to L. Shan).

## Author Contributions

L.S. and Q.W. designed the study, analyzed the data and wrote the manuscript; Q.W. performed the experiments; P.T. and C.D. conducted microscopy experiments; S.K. contributed to immunoblotting analysis of protease functions; R.P. supervised the studies using patient samples; H.G. contributed to generation of viral constructs; G.H. processed patient and healthy donor blood samples; K.C. analyzed the data on infection with clinical HIV-1 isolates.

## Competing interests

The authors declare no competing financial interests.

**Table S1.**
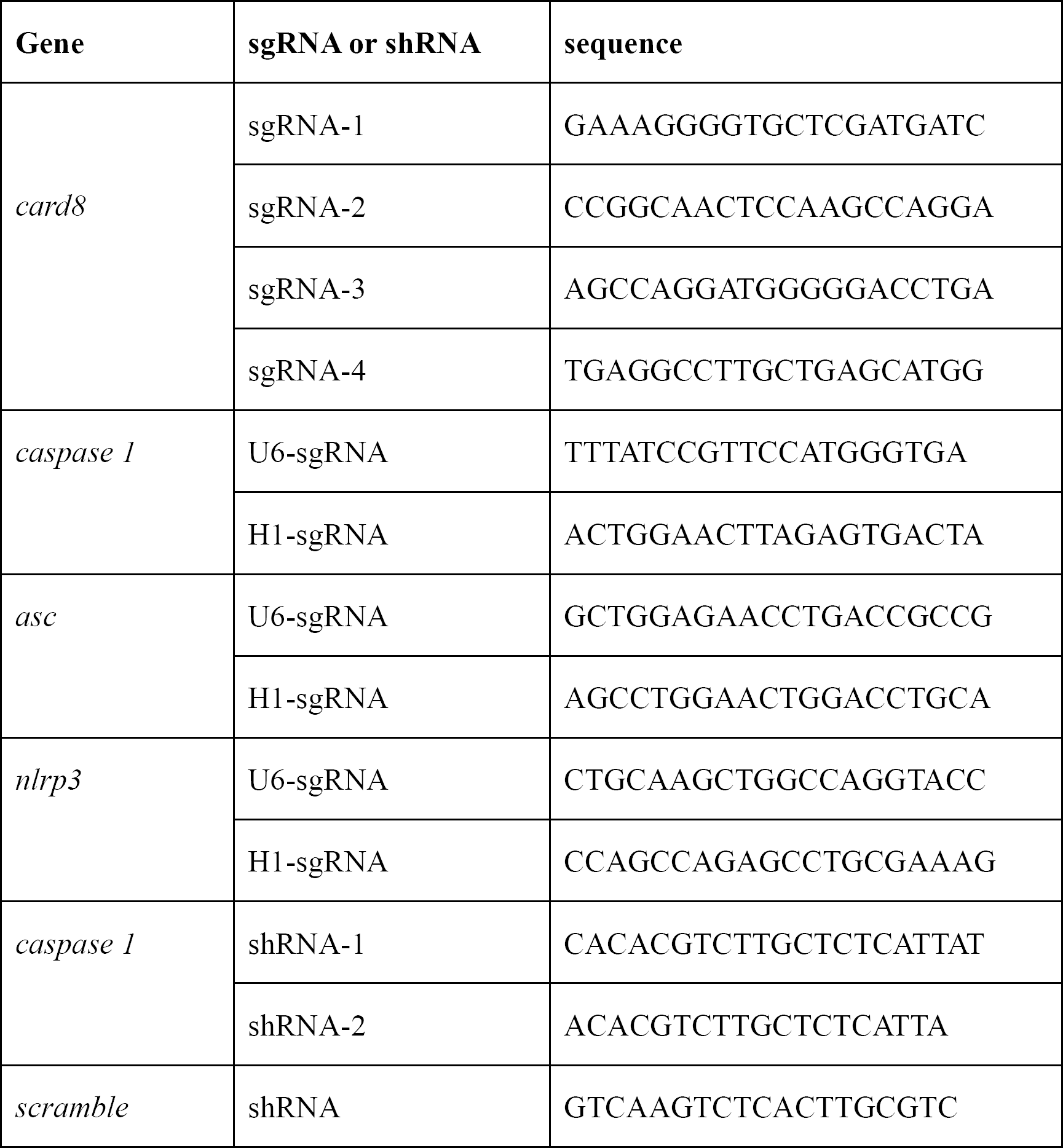
sgRNA and shRNA sequences.

**Figure S1:**
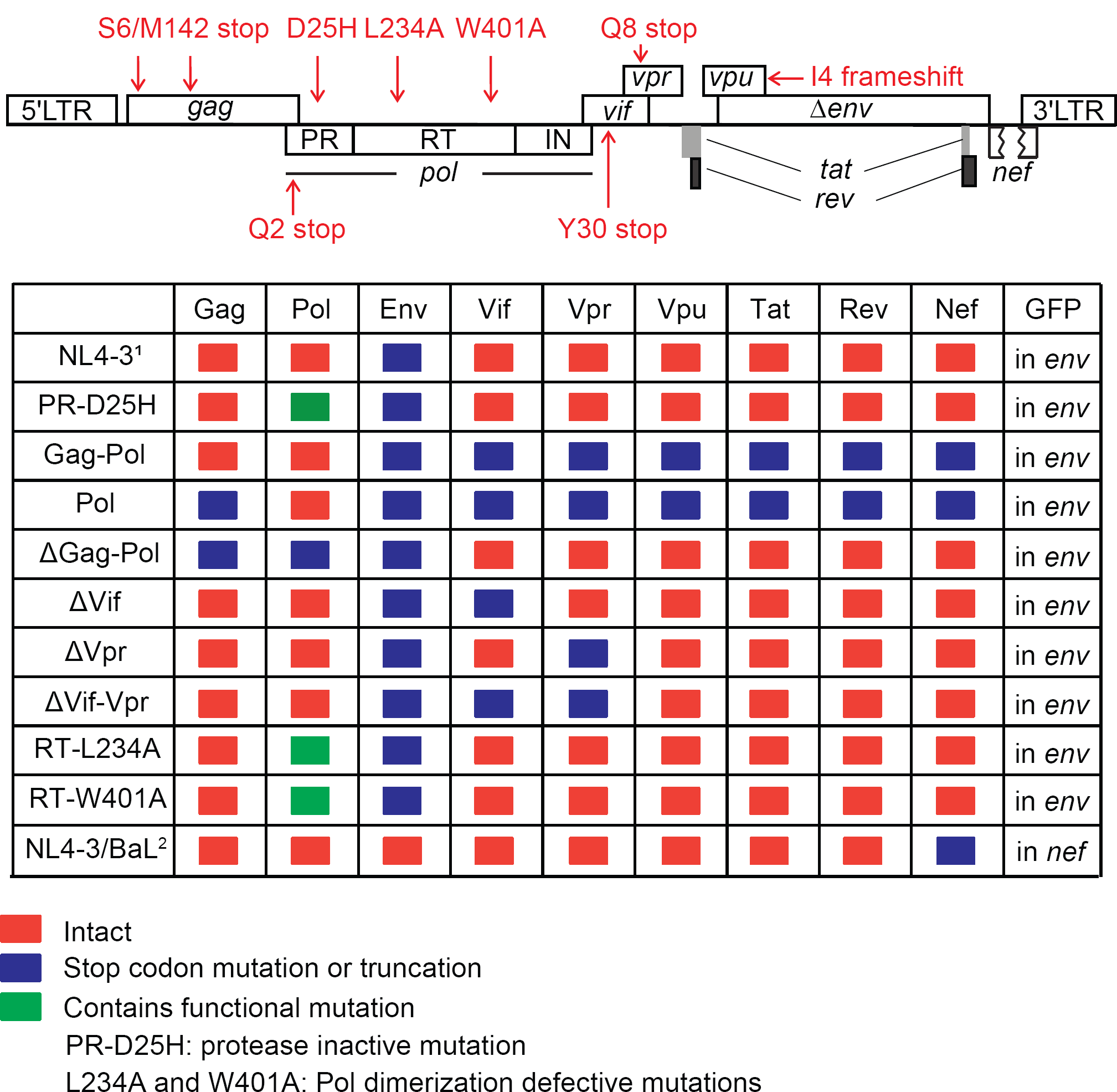
A list of stop codon mutations or truncations. ^1^: All viral plasmids were generated using the NL4-3-ΔEnv-EGFP backbone, except NL4-3/BaL. ^2^: NL4-3/BaL has the NL4-3 backbone with EGFP coding sequencing in *nef*, and the BaL envelop coding sequence. These plasmids were used to transfect HEK293T cells, either for immunoblotting analysis of CARD8 and IL-1β cleavage, or for production of HIV-1 reporter viruses when co-transfected with pVSV-G and lentivirus packaging plasmid. HIV-1 reporter viruses generated using NL4-3-ΔEnv-EGFP backbone are replication-defective. The viral vector NL4-3/BaL can produce replication-competent viruses.

**Figure S2:**
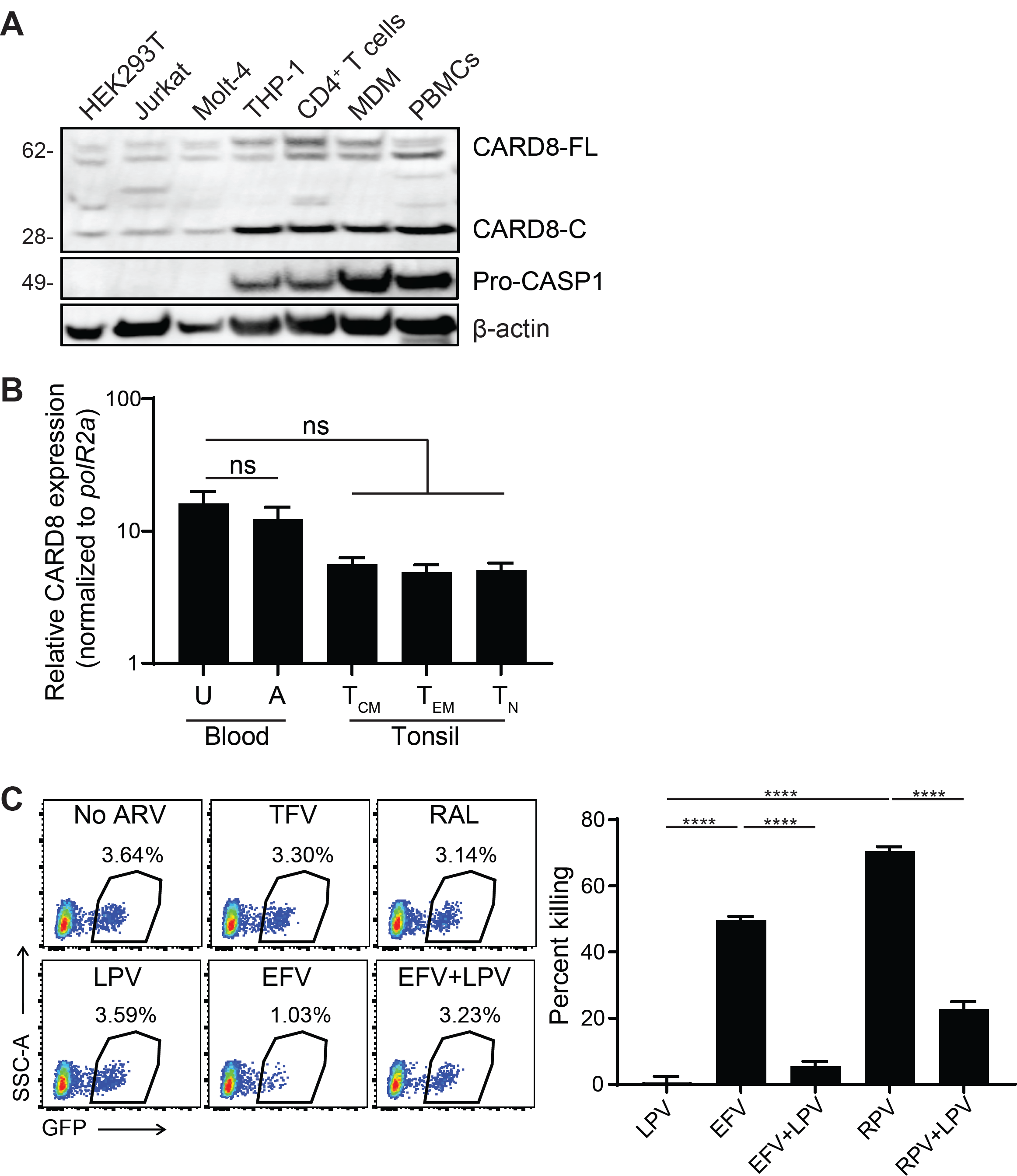
Expression of CARD8 and killing of unstimulated CD4^+^ T cells. **(A)** CARD8 expression in cell lines, as well as primary CD4^+^ T cells, monocytes derived macrophages and total PBMCs. Cell lysates were evaluated by immunoblotting using antibodies against CARD8 C-terminus or CASP1. CARD8-FL, full-length CARD8; CARD8-C, C terminal CARD8. **(B)** CARD8 transcription in primary CD4^+^ T cells. Central memory (T_CM_, CD45RO^+^CCR7^+^), effector memory (T_EM_, CD45RO^+^CCR7^-^) and naïve (T_N_, CD45RO^-^CCR7^+^) CD4^+^ T cells were purified from the CD3^+^CD4^+^CD8^-^ fraction of tonsil mononuclear cells by sorting. Blood CD4^+^ T cells were isolated from healthy donor PBMCs. U: unstimulated; A: activated. CARD8 transcription was measured by qPCR. **(C)**, Unstimulated CD4^+^ T cells were infected with HIV-1 reporter virus NL4-3-Pol. 4 days post infection, cells were treated with indicated ARVs for 24 hours before FACS analysis. *P* values were calculated using one-way ANOVA and Turkey multiple comparison tests. ****p < 0.0001. N=3. Error bars show mean values with SEM.

**Figure S3:**
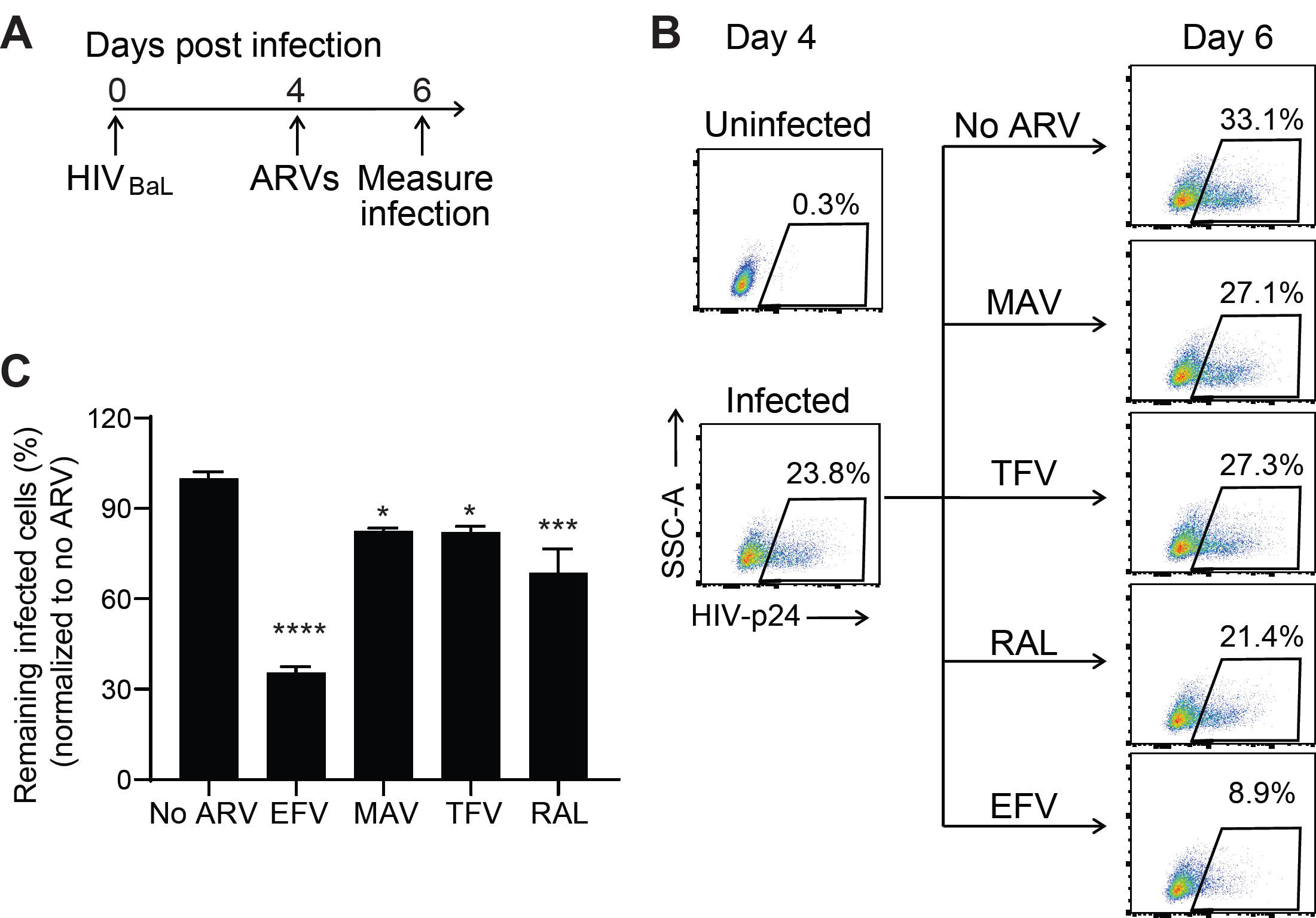
Killing of primary CD4^+^ T cells infected with HIV_BaL_. Activated primary CD4^+^ T cells were infected with HIV_BaL_. At day 4, ARVs were added at 5μM. Intracellular p24 staining was performed to measure frequency of HIV-1-infected cells at day 6. Infection was normalized to no ARV control. *P* values were calculated using one-way ANOVA and Turkey multiple comparison tests. *p < 0.05; ***p < 0.001; ****p < 0.0001. N=3. Error bars show mean values with SEM.

**Figure S4:**
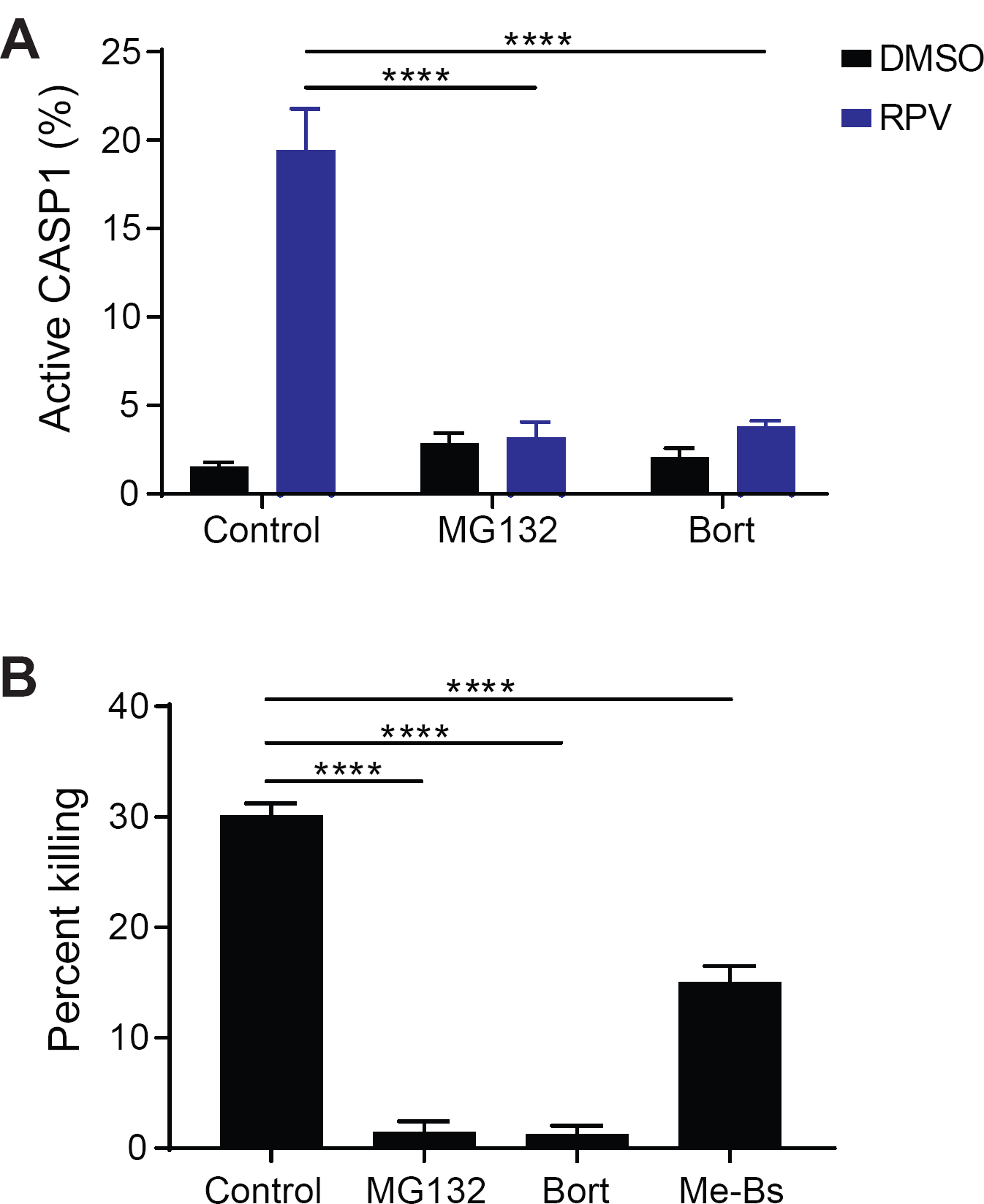
Proteasome inhibitors block HIV-1 protease-mediated inflammasome activation in HIV-1-infected primary CD4^+^ T cells. Activated primary CD4^+^ T cells were infected with VSV-G pseudotyped HIV-1 reporter virus NL4-3-Pol. At day 3 post infection, cells were pre-treated with MG132 (10μM), Bort (5μM), or Me-Bs (20μM) for 30 minutes and then treated with RPV for 6 hours. In **A**, FLICA660 Caspase 1 was used to measure active CASP1. In **B**, cell killing was determined by FACS. In **A**, *P* values were calculated using two-way ANOVA and Turkey multiple comparison tests. In **B**, *P* values were calculated using one-way ANOVA and Turkey multiple comparison tests. ****p < 0.0001. In each bar graph, n=3. Error bars show mean values with SEM.

**Figure S5:**
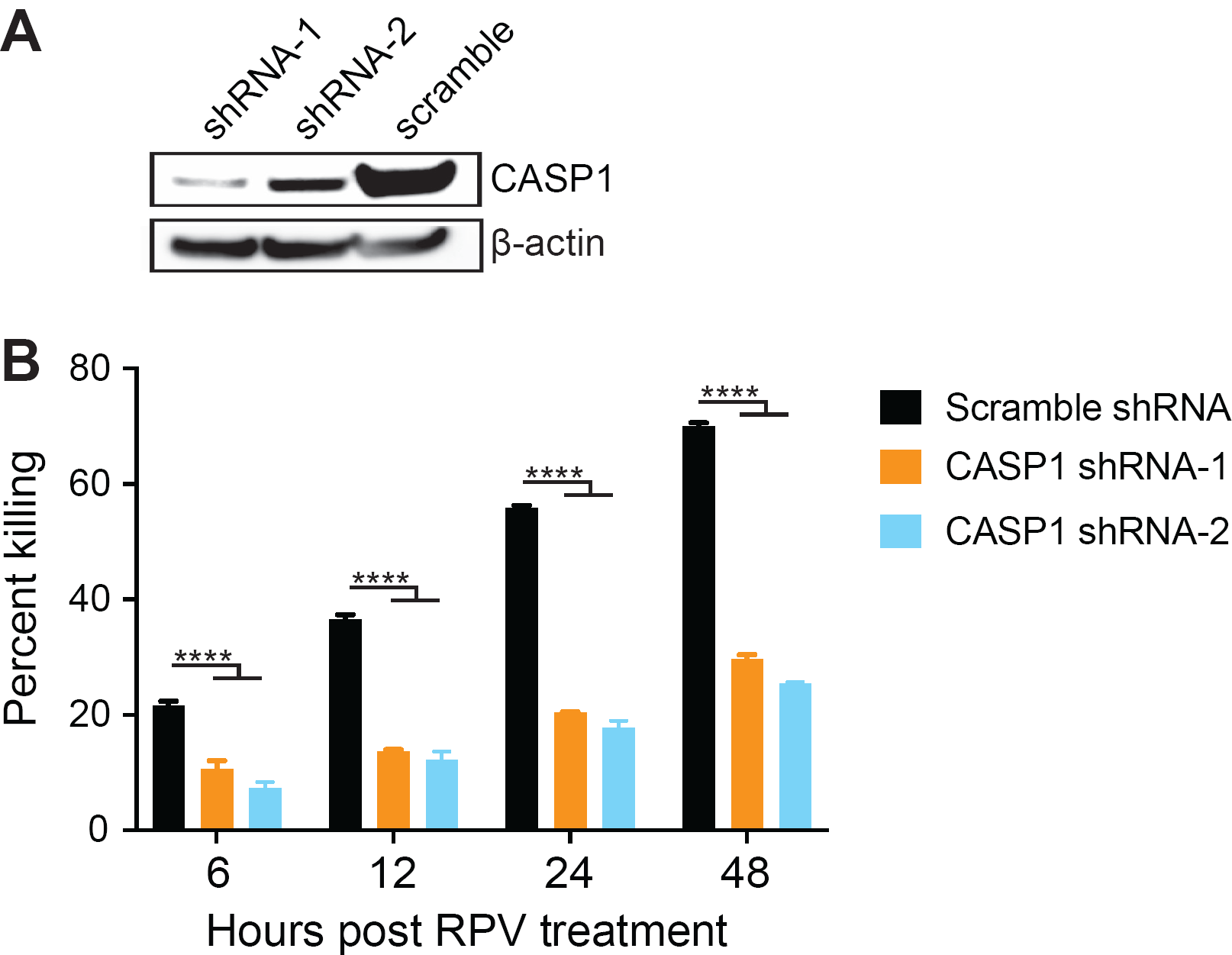
Killing of HIV-1-infected primary CD4^+^ T cells is blocked by knockdown of CASP1. **(A)** Validation of CASP1 knockdown in primary CD4^+^ T cells by immunoblotting. **(B)** CASP1-KD primary CD4^+^ T cells were infected with VSV-G pseudotyped HIV-1 reporter virus NL4-3-Pol. 3 days after infection, cells were treated with RPV. GFP expression was analyzed by FACS 24 hours post RPV treatment. *P* values were calculated using two-way ANOVA and Turkey multiple comparison tests. ****p < 0.0001. N=3. Error bars show mean values with SEM.

**Figure S6:**
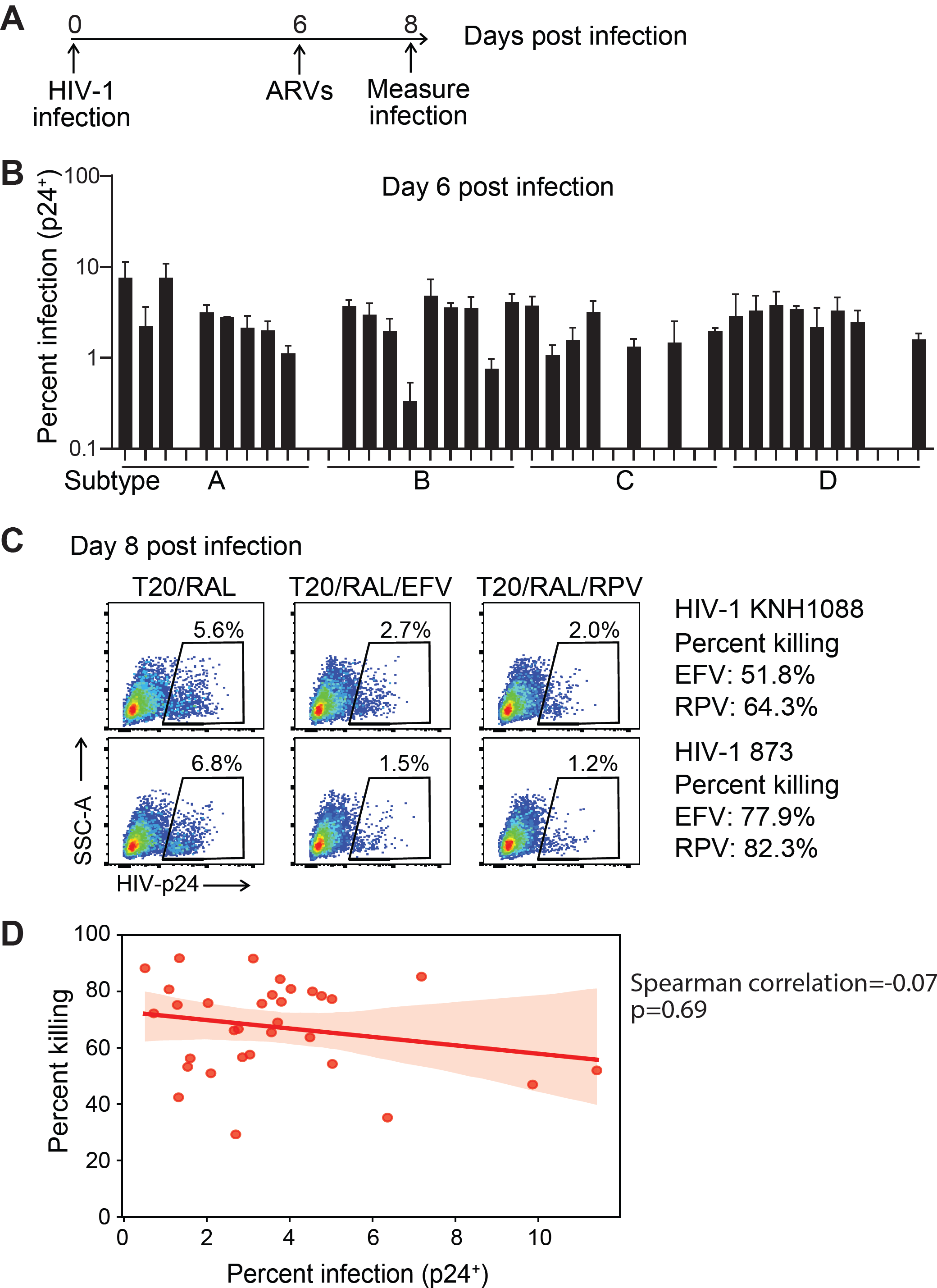
Killing of primary CD4^+^ T cells infected with clinical HIV-1 isolates. **(A)** Scheme of the infection experiments. Primary CD4^+^ T cells were infected with clinical HIV-1 isolates. RAL (5μM) and T-20 (1μM) with or without EFV or RPV (5μM) were added on day 6 to block new infection. The remaining infected cells were measured at day 8. **(B)** Infection with clinical HIV-1 isolates. Activated primary CD4^+^ T cells were infected with clinical isolates. Intracellular p24 staining was performed to determine the frequency of HIV-1 infected at day 6. Infection was performed with three independent healthy donor CD4^+^ T cells and the average values were shown. N=3. Error bars show mean values with SEM. HIV-1 isolates with <0.3% infection were excluded from killing analysis in the Figure 4a. **(C)** Killing of two representative clinical isolates HIV-1 KNH1088 (subtype A) and HIV-1 873 (subtype B). **(D)** Correlation between viral fitness (frequency of p24^+^ cells in T20/RAL group at day 8) and killing efficiency of EFV. For correlation analysis, infection data from one healthy donor were used. The percent killing and percent infection were imported into python (v.3.7.3 ref.1) using the package pandas (v.1.0.1 ref.2). The data was visualized using the seaborn (v.0.10.0 ref.3) and matplotlib packages (v.3.1.3 ref.4) and a spearman correlation was done using the scipy stats module pearmnr (v 1.4.1 ref.5) to obtain correlation coefficients and p-values.

**Figure S7:**
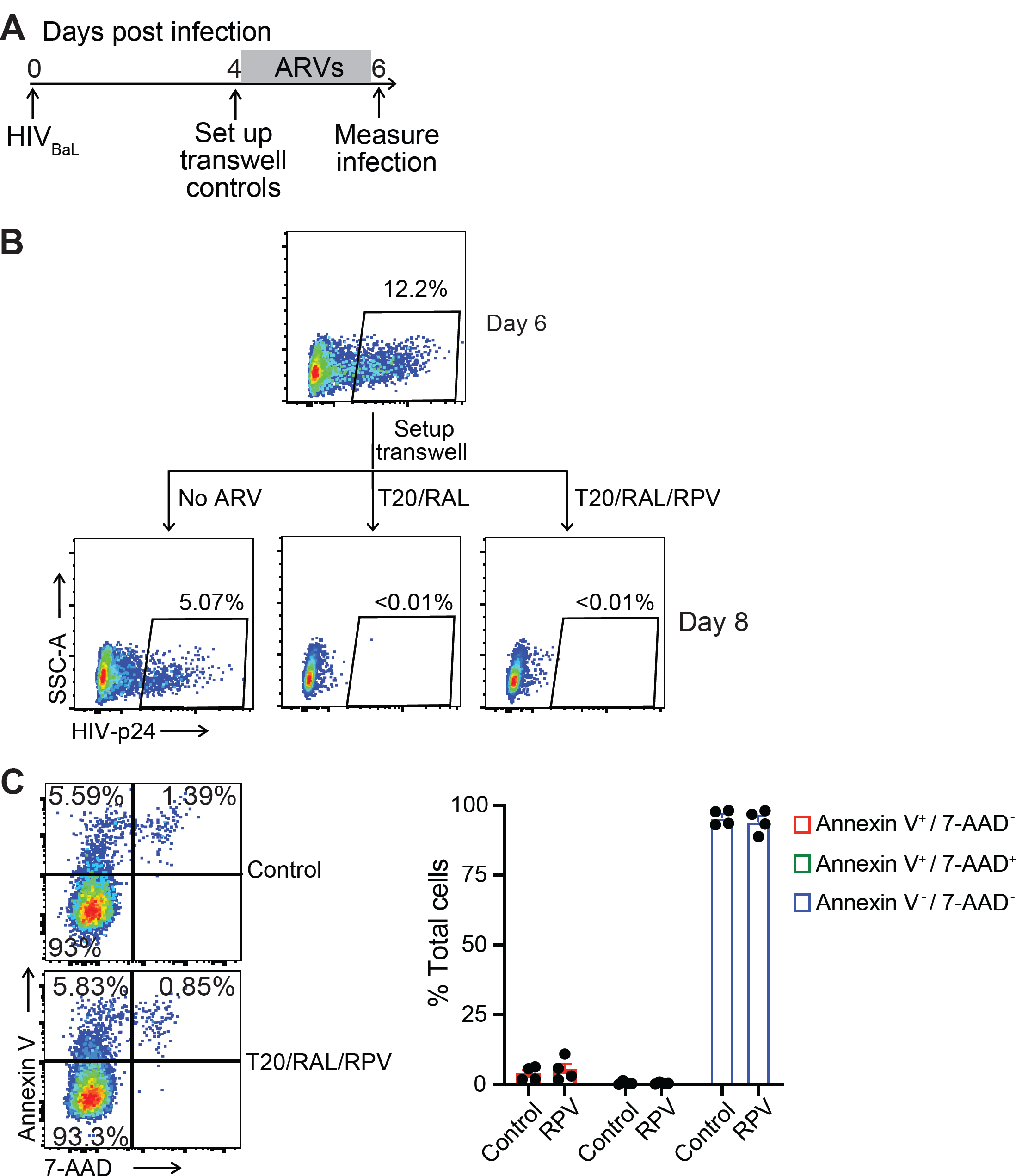
Viral suppression and cytotoxicity of ARV combinations. **(A-B)** ARV combinations are sufficient to prevent new rounds of HIV-1 infection. Activated primary CD4^+^ T cells were infected with HIV_BaL_. At day 4 post infection, cells were seeded into the upper layer of transwell plates. Uninfected cells were seeded into the bottom layer of transwell plates. RAL (5μM) and T-20 (1μM) with or without RPV (5μM) were added. Cells from bottom layer were collected on day 6 for intracellular p24 staining. **(C)** ARV combinations are not cytotoxic. Activated primary CD4^+^ T cells from HIV-1 patients were treated with RAL (10μM) and T-20 (10μM) with or without RPV (5μM) for 48 hours. Cell viability analysis was performed by Annexin V and 7-AAD staining. Experiments were performed using 4 independent blood samples.

**Video S1: pyroptosis of HIV-1-infection macrophages**.

Viral infection and drug treatment were described in Figure 2D. HIV-1-infected macrophages were treated with DMSO or RPV (5μM). Imaging began immediately after drug treatment. Images were acquired at 3 minutes intervals for 6 hours. Images were processed and stitched using Gen5 software (Biotek). Representative time-lapse video depicts a single field over 6 hours.

